# Selecting genomes that matter: haplotype-based prioritization for iterative pangenome expansion

**DOI:** 10.64898/2026.05.13.724976

**Authors:** Marina P. Marone, Erwang Chen, Axel Himmelbach, Georg Haberer, Manuel Spannagl, Nils Stein, Martin Mascher

## Abstract

**Background:** As pangenomes approach saturation, identifying additional genomes that contribute novel sequence information becomes increasingly difficult. Current sample-selection strategies often rely on global diversity metrics or variant counts and do not explicitly account for the composition of an existing pangenome, a limitation that becomes increasingly relevant as pangenomes mature. Here, we present SelHap, a haplotype-based pipeline that uses whole-genome sequencing (WGS) data to prioritize accessions based on their contribution of novel haplotypes relative to a defined background, enabling targeted and iterative pangenome expansion.

**Results:** We applied SelHap to the barley pangenome, using 76 assembled genomes as a background to select new accessions from a large WGS panel. Using this approach, we generated chromosome-scale genome assemblies from 19 accessions selected with SelHap and from 17 elite lines selected based on their relevance in historical barley breeding. Across multiple benchmarking scenarios, SelHap-based selection consistently resulted in a greater increase in non-redundant (single-copy) pangenome sequence, demonstrating that prioritizing haplotype novelty relative to an existing background maximizes unrepresented sequence content.

**Conclusions:** By transforming complex haplotype-clustering outputs into interpretable summaries and ranked candidate lists, SelHap provides a practical framework for targeted pangenome expansion. Beyond sample selection, SelHap can facilitate ancestry and germplasm comparisons across diverse panels. As WGS data become more accessible, SelHap offers a scalable and interpretable solution for extending mature pangenomes by explicitly targeting previously unrepresented sequence space.

## Introduction

As plant pangenomes mature, selecting additional genomes that contribute genuinely new information has become increasingly challenging. Sample selection remains an essential step, not only to ensure adequate representation of genetic diversity, but also to tailor it to stakeholders needs such as breeders. Popular cultivars and useful lines are often included, yet other accessions can be selected to maximize representation of genetic diversity. Sample selection based on distinct geographic origins, taxonomic classifications or morphological traits can provide a distinct enough dataset, but more informed decisions can be made by using sequencing data.

Currently, the most widely used sequencing strategies for diversity analyses are GBS (genotyping by sequencing) and WGS (whole-genome shotgun sequencing), which mainly differ in marker density. With such marker datasets, it is possible to calculate genetic distances and obtain similarity metrics like identity-by-state (IBS) or to summarize genomic data structure using principal component analysis (PCA). In addition, specialized tools also use marker sets for sample selection, such as Core Hunter 3 (1), which maximizes genetic diversity, and SVCollector (2), which prioritizes highest number of structural variants (SVs). Although these approaches are useful for initial sample selections, they do not account for previously represented genetic diversity nor for haplotype-level information. As datasets grow and pangenomes approach saturation, tools capable of incorporating broader biological signals to guide strategic sample selection are increasingly needed.

To address this gap, we developed SelHap, a WGS-based pipeline to select samples that carry haplotypes not yet represented in a pangenome. The pipeline compares a broad panel to a user-defined background, both present in a marker dataset. SelHap offers three running modes: i) selecting long haplotype stretches absent from the background, ii) applying the same principle while considering haplotype frequency, or iii) targeting specific regions. We applied SelHap to the barley pangenome, which currently contains 76 high-quality chromosome-scale genome assemblies, most selected using PCA. Our goal was to include additional cultivars relevant to the Central European breeding germplasm, as well as haplotypes not yet explored in the existing pangenome. Our central hypothesis is that prioritizing accessions based on haplotype novelty relative to an existing pangenome maximizes the addition of previously unrepresented sequence space more effectively than strategies based solely on global diversity summaries.

In this paper, we describe the SelHap pipeline and evaluate its utility by applying it to the selection of new accessions for the barley pangenome. Through this approach, we identified 19 barley genotypes carrying previously unrepresented haplotypes. To capture key germplasm used in European breeding programs, we manually selected 17 elite barley cultivars. All 36 genotypes were sequenced with high-fidelity (HiFi) long-reads (PacBio Revio) and Hi-C, enabling the construction of chromosome-scale genome assemblies.

## Results

### Overview of the SelHap pipeline

SelHap is a pipeline for selecting new accessions to integrate a pangenome. It builds on the output of IntroBlocker (3), a tool that assigns ancestral haplotype groups (AHGs) to genomic bins, to investigate fine-scale ancestry and evolutionary history. SelHap addresses a simpler but distinct problem. It uses these haplotype clusters to optimize haplotype novelty relative to an existing pangenome, fundamentally differing from approaches that maximize global diversity (PCA, Core Hunter 3) or SV counts (SVCollector).

The pipeline requires WGS reads - or simulated reads from a genome assembly - from the pangenome accessions and the candidate samples. Reads are aligned to a reference of choice to then call the variants, obtaining a VCF file. After filtering with adequate parameters, this VCF file will be used as input for IntroBlocker to obtain haplotype clusters. Although IntroBlocker produces per-chromosome plots of haplotype blocks, varying sample order across chromosomes makes large datasets difficult to interpret directly. These plots are created from tables that record clustered haplotypes per genomic window. SelHap operates on these tables to identify haplotypes absent from the background (existing pangenome), but present in the foreground (candidate accessions). By doing so, it highlights accessions that contribute new haplotypes to the pangenome (**Fig. 1A**).

**Figure 1:**
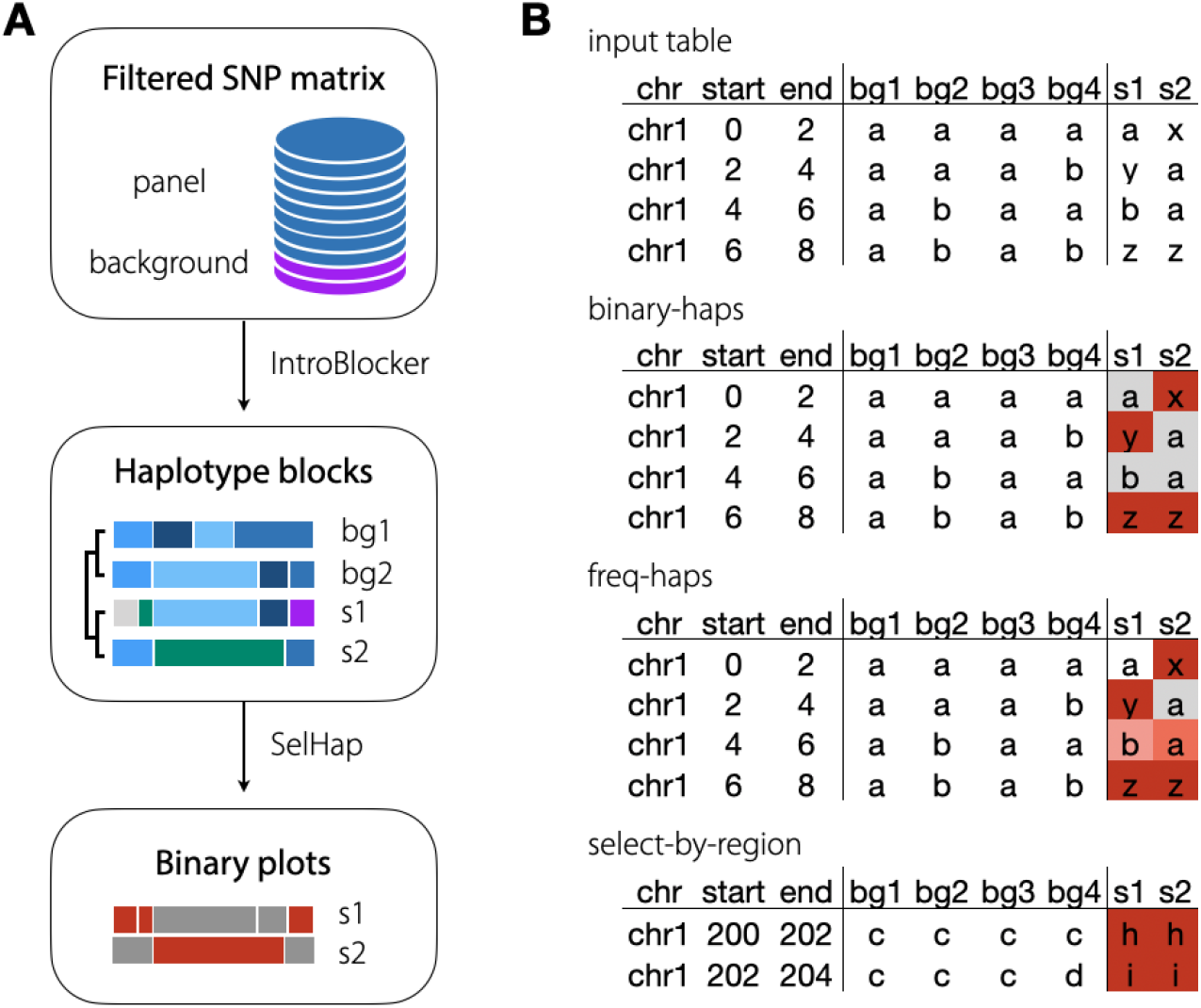
Overview of the SelHap pipeline. **A:** Graphical summary of the steps needed to run SelHap. A filtered SNP matrix with background samples (already existing pangenome dataset) and putative samples is input into IntroBlocker, which generates groups of clustered haplotype blocks per chromosome, and whose output tables will be used as input to SelHap. The foreground samples will be compared to the background samples to check if the haplotype on each window is new in relation to the background. If it is new, it is colored red; otherwise, gray. **B:** Examples of how the comparisons take place according to each running mode. Binary-haps check for presence or absence only, while freq-haps gives a score to the frequency each haplotype appears on the background. Select-by-region will do the same as binary-haps, but for specific ranges on the chromosomes. bg1-bg4: background samples. s1 and s2: samples being tested.

SelHap has three running modes: binary-haps, freq-haps, and select-by-region (**Fig. 1B**). In all three modes, the output consists of tables with scores per chromosome for all samples and, optionally, per-chromosome plots with each window of haplotypes colored by their presence (gray), absence (red) or frequencies. The base running mode is binary-haps, that will check if a specific genomic window haplotype is present or absent in the background. Therefore, if no sample in the background has the tested haplotype in that genomic window, this sample’s haplotype is assigned as novel in relation to the background. To account for long novel haplotypes, we apply run-length encoding (RLE) in R and square the resulting segment lengths to weigh longer haplotypes more strongly, so samples with longer novel haplotypes receive a higher score. This process is done independently for each chromosome and at the end chromosome-specific scores are summed to obtain a genome-wide score for each sample. Samples are then ranked by decreasing score and the top-scoring one is added to the background set so the process is repeated for a number of user-defined iterations.

The other two modes are variations of binary-haps. Frequency-haps works the same way, but it gives a rating from 1 to 10 according to the frequency of the tested haplotype. The higher the score, the more abundant this haplotype is in the background. Another potential application is the comparison of different germplasm groups to assess haplotype origins, relatedness, and patterns of ancestry. Lastly, select-by-region will use results of 1-iteration binary-haps to select samples with new haplotypes in a user-specified genomic region. The proportion of new haplotypes in the range can be set by the user.

### Applying SelHap to the barley pangenome

We evaluated SelHap on a large barley panel consisting of 1,769 genotypes with available WGS data and simulated reads for the 76 barley pangenome version 2 (BPGv2) assemblies (**Sup. Table S1**). Across all 1,845 accessions, we identified over 24 million SNPs, which provided dense marker coverage for haplotype clustering. We ran IntroBlocker with this dataset, using the semi-supervised algorithm with a threshold of 500 SNPs per 2 Mb window. The output tables were used to run SelHap in binary-haps mode to obtain a total of 20 samples with novel haplotypes compared to the 76 pangenome accessions as the background.

Running SelHap binary-haps revealed a strong enrichment of novel haplotypes in wild barley. The top-ranked accessions comprise two landraces and 18 wild samples (**Sup. Table S2, Sup. Fig. S1**), showing long stretches of haplotypes absent from the 76 pangenome accessions, particularly in the distal regions. This pattern was expected, given the limited representation of wild samples in the current pangenome and the unexplored diversity present in wild relatives (4-6). Nevertheless, the top ranked accession was HOR 18876, a Japanese landrace. The runtime on a laptop (MacBook Air, 2022 model with Apple M2 chip, 8-core CPU, 8-core GPU, 16 GB unified memory) for 1 iteration was 27 seconds, and to iterate through 19 more samples, 8:52 minutes, demonstrating that the pipeline is fast even when applied to large datasets.

Since our focus was on haplotypes closer to the breeding germplasm, we removed wild samples and repeated this analysis. The refined list contained 8 cultivars and 12 landraces (**Table 1, Sup. Table S2, Sup. Fig. S2**), with HOR 18876 consistently appearing at the top, due to its long novel haplotype segments on all chromosomes. Runtime with 1,602 samples still remained comparable (23 seconds and 7:31 minutes to iterate to 20 samples). From these 20, we selected 15 accessions to integrate to the barley pangenome after verifying seed availability (**Table 1**).

**Table 1:** SelHap output for a run with the barley pangenome accessions as background, excluding wild samples from the foreground. This was run with the binary-haps iterative mode up to 20 samples. The samples in bold were included in the first test dataset, while the ones with an asterisk were included in the second.

| Sample | chr1H | chr2H | chr3H | chr4H | chr5H | chr6H | chr7H | Sum |
| --- | --- | --- | --- | --- | --- | --- | --- | --- |
| **HOR 18876** | 704 | **9937** | 934 | **9301** | 5977 | 2786 | **4422** | 34061 |
| **HOR 15692** | **7069** | 138 | 4668 | 40 | 52 | **4329** | 41 | 16337 |
| **HOR 15254 (Smooth Awn 86)** | 173 | 50 | 730 | **4066** | 286 | **2941** | **2455** | 10701 |
| **Balder J \*** | 223 | 454 | **4317** | 160 | **2537** | **2207** | 315 | 10213 |
| **HOR 20348** | 69 | **2201** | 110 | 735 | **3528** | **2035** | 184 | 8862 |
| **HOR 14307** | 39 | 67 | **8177** | 120 | 358 | 20 | 13 | 8794 |
| Andreia (ACK-16) | 141 | 198 | 512 | 907 | **5444** | 671 | 286 | 8159 |
| **Proctor \*** | 28 | 845 | **3806** | 44 | **1498** | 590 | 557 | 7368 |
| **HOR 18382** | 71 | **6648** | 197 | 21 | 109 | 79 | 44 | 7169 |
| **HOR 2805** | 22 | 389 | 50 | 12 | 34 | **5466** | **1106** | 7079 |
| **Okal** | 28 | 8 | 39 | 0 | 159 | **347** | **5684** | 6265 |
| **HOR 1062** | 13 | 10 | 6 | 404 | 455 | **4878** | 201 | 5967 |
| **HOR 16149** | 311 | 201 | 63 | 402 | 198 | 11 | **4747** | 5933 |
| **HOR 11419** | 20 | 456 | 47 | 345 | **4668** | 77 | 168 | 5781 |
| **Spey \*** | 159 | **1289** | 399 | 1082 | 1075 | 934 | 804 | 5742 |
| HOR 17538 | 18 | 18 | **5436** | 8 | 112 | 8 | 42 | 5642 |
| **Rika \*** | 2 | 18 | **4314** | 54 | **1036** | 6 | 167 | 5597 |
| Arthur | 45 | 338 | **4208** | 7 | 20 | **445** | 131 | 5194 |
| Focus * | 18 | 16 | **4700** | 6 | 35 | 176 | 5 | 4956 |
| HOR 17386 | 203 | 500 | 62 | **1624** | 195 | **1701** | 182 | 4467 |

We next checked whether these accessions carried novel haplotypes in pericentromeric regions, which were previously reported to be largely saturated in the barley pangenome dataset (7). Using SelHap’s select-by-region mode and requiring at least 50% of novel haplotypes within ± 80 Mb of the Morex centromeres, we identified 34 accessions with novel haplotypes on at least one of the chromosomes (**Sup. Table S3, Sup. Fig S2**). Nearly all accessions (19 out of 20) overlapped with those ranked highly in the iterative genome-wide analysis. Additionally, we selected 4 spring 2-rowed cultivars from the SelHap rankings that showed clear pericentromeric novelty (**Sup. Table S3**). We plotted all final 19 samples to illustrate the conceptual differences between bin-haps and freq-haps modes (**Fig. 2**).

**Figure 2:**
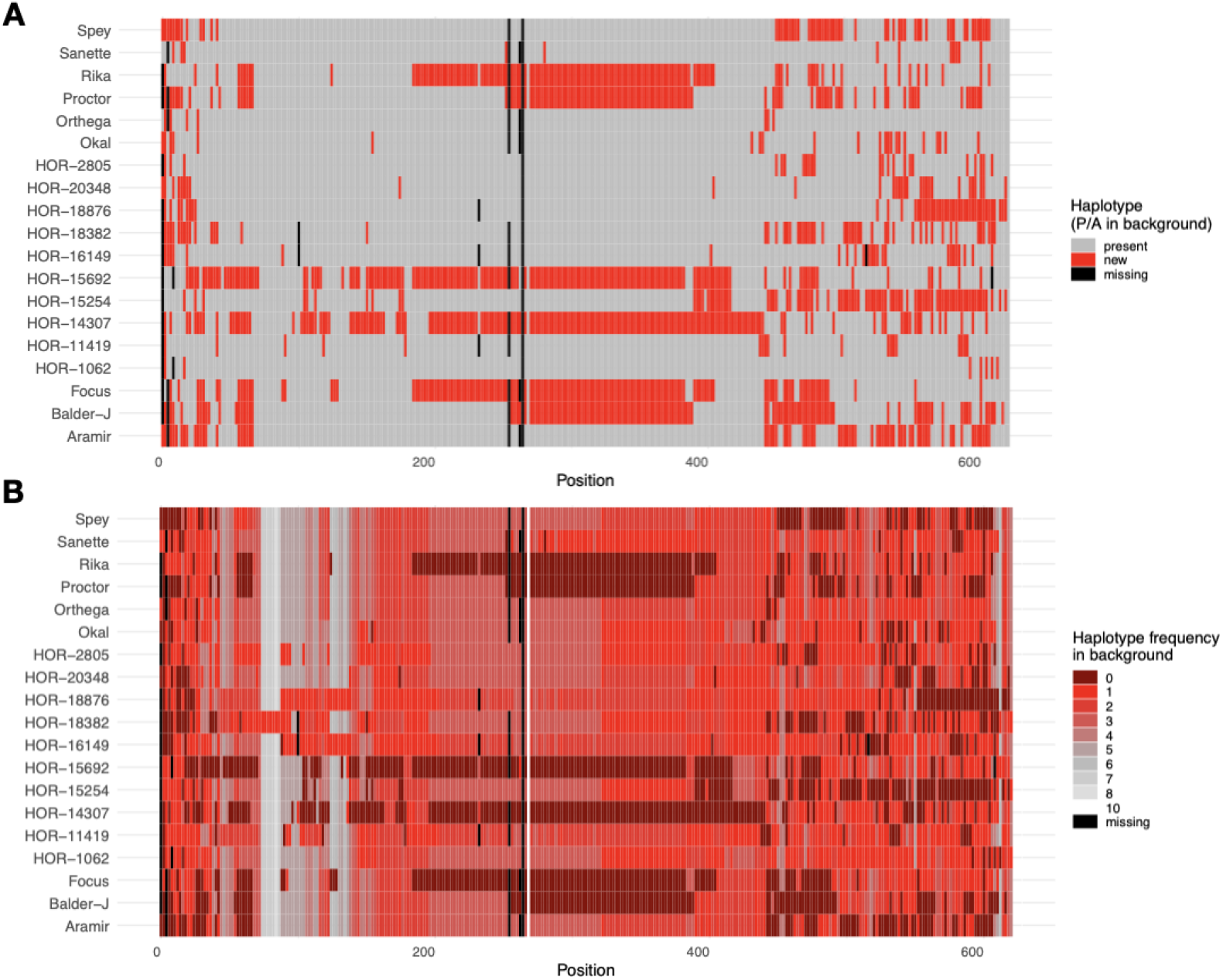
Example output plots for barley chromosome 3H of the 19 samples selected with SelHap. **A:** Binary-haps output plot from 1 iteration. Gray positions show haplotypes present in the background (BPGv2), while red show novel haplotypes and black show missing data. **B:** Freq-haps output plot. The color scale depicts the haplotype frequency ranks computed from the background dataset, where lower ranks indicate less frequent haplotypes, being the ones in dark red (0) completely absent in the background. Genomic windows have 2 Mb.

To assess SelHap’s robustness, we repeated the analysis using more stringent IntroBlocker parameters. We ran binary-haps with 20 iterations for each dataset, increasing either SNP threshold (500 per window) or window size (5 Mb). In both cases, fewer novel haplotype stretches were detected, consistent with stricter filtering (**Sup Fig. S4A**). Nevertheless, sample selection remained relatively stable. Increasing SNP threshold resulted in 90% (18/20) overlap, suggesting that the main haplotype signal is preserved while reducing background noise. In contrast, increasing the window size resulted in lower overlap (70%, 14/20), likely because larger windows redefine haplotype blocks and alter contributions of individual regions to the overall score (**Sup. Fig. S4B**).

### Genome assemblies of 36 barley cultivars and landraces

To obtain a dataset that would complement the barley pangenome with more cultivars and novel haplotypes, we assembled genomes for a total of 36 accessions: 19 selected with SelHap and 17 elite cultivars chosen by barley breeders or through pedigree records (**Table 2**). Genomes were assembled with PacBio HiFi and 3D conformation capture sequencing (Hi-C) data, achieving high-quality chromosome-scale assemblies. The contig N50 values ranged from 4.9 to 16 Mb, and the total genome sizes varied between 4.19 and 4.30 Gb, consistent with the expected barley genome size. BUSCO analysis indicated >97.2% completeness across all assemblies, confirming their high accuracy and integrity (**Sup. Table S4 and S5**).

**Table 2:** Samples sequenced in this study using different selection criteria.

| Name | Method | Status | Growth habit | Row number | Country | Release year | Assembly size (Gb) |
| --- | --- | --- | --- | --- | --- | --- | --- |
| Valticky | breeders | cultivar | S | 2 | CSK | 1938 | 4.21 |
| Scarlett | breeders | cultivar | S | 2 | DEU | 1995 | 4.23 |
| Quench | breeders | cultivar | S | 2 | GBR | 2006 | 4.23 |
| Diamant | breeders | cultivar | S | 2 | CSK | 1965 | 4.20 |
| Annabell | breeders | cultivar | S | 2 | DEU | 1999 | 4.22 |
| Saffron | breeders | cultivar | W | 2 | GBR | 2003 | 4.22 |
| Retriever | breeders | cultivar | W | 2 | GBR | 2007 | 4.22 |
| Maris Otter | breeders | cultivar | W | 2 | GBR | 1965 | 4.19 |
| Vogelsanger Gold | breeders | cultivar | W | 6 | DEU | 1965 | 4.22 |
| Theresa | breeders | cultivar | W | 6 | DEU | 1994 | 4.22 |
| Franka | breeders | cultivar | W | 6 | DEU | 1980 | 4.19 |
| Esterel | breeders | cultivar | W | 6 | FRA | 1995 | 4.23 |
| Triumph | pedigree table | cultivar | S | 2 | DEU | 1965 | 4.22 |
| Maythorpe | pedigree table | cultivar | S | 2 | GBR | 1954 | 4.20 |
| Heines Haisa I | pedigree table | cultivar | S | 2 | DEU | 1939 | 4.20 |
| Hanna | pedigree table | cultivar | S | 2 | CSK | 1884 | 4.21 |
| Malta | pedigree table | cultivar | W | 2 | DEU | 1968 | 4.20 |
| Spey | SelHap | cultivar | S | 2 | GBR | 1995 | 4.24 |
| Sanette | SelHap | cultivar | S | 2 | DNK | 2012 | 4.22 |
| Rika | SelHap | cultivar | S | 2 | SWE | 1949 | 4.20 |
| Proctor | SelHap | cultivar | S | 2 | GBR | 1952 | 4.30 |
| Orthega | SelHap | cultivar | S | 2 | CSK | 1998 | 4.30 |
| Focus | SelHap | cultivar | S | 2 | DEU | 2018 | 4.24 |
| Balder J | SelHap | cultivar | S | 2 | SWE | 1964 | 4.21 |
| Aramir | SelHap | cultivar | S | 2 | NLD | 1972 | 4.21 |
| Smooth Awn 86<br>(HOR 15254) | SelHap | cultivar | W | 6 | USA | 1939 | 4.26 |
| Okal | SelHap | cultivar | W | 6 | SVK | 1992 | 4.23 |
| HOR 11419 | SelHap | landrace | S | 6 | IND | NA | 4.26 |
| HOR 14307 | SelHap | landrace | S | 6 | AFG | NA | 4.20 |
| HOR 2805 | SelHap | landrace | S | 6 | IRN | NA | 4.30 |
| HOR 15692 | SelHap | landrace | S | NA | AFG | NA | 4.30 |
| HOR 20348 | SelHap | landrace | S | NA | ETH | NA | 4.23 |
| HOR 18382 | SelHap | landrace | S | NA | PAK | NA | 4.22 |
| HOR 18876 | SelHap | landrace | S | NA | JPN | NA | 4.30 |
| HOR 1062 | SelHap | landrace | W | 6 | TUR | NA | 4.24 |
| HOR 16149 | SelHap | landrace | W | NA | TUR | NA | 4.24 |

Gene annotation based on projection from previous barley pangenome assemblies (7) identified, on average, a total of 41,575 genes (σ = ±166) in the 36 accessions. Of these, 40,028 (σ = ±162), 473 (σ = ±4.4) and 1,074 (σ = ±16) genes were classified as protein-coding, plastid- and transposon-related, respectively (**Sup. Table S5; Sup. Fig. S5**). For the 36 accessions, we determined on average 4812 (σ = ±12), 13 (σ = ±12), 71 (σ = ±11), complete, fragmented and missing BUSCO genes, respectively (**Sup. Table S5; Sup. Fig. S6**).

### Comparison to cultivar dataset curated by breeders

A direct comparison between haplotype-based sample selection and cultivar sets curated by breeders requires careful interpretation because the two strategies optimize different objectives. Breeder selection is primarily driven by agronomic performance, adaptation and pedigree considerations rather than by the maximization of genomic novelty. Consequently, differences in germplasm composition, including the relative contribution of cultivars and landraces, may influence comparisons of diversity and structural variation. Nevertheless, breeder-curated cultivar sets provide a practically relevant benchmark because they reflect realistic accession-selection decisions in applied barley genomics.

To assess whether prioritizing haplotype novelty relative to an existing pangenome translates into increased genomic novelty in assembled genomes, we compared accessions selected with SelHap to a set of cultivars curated by breeders (**Sup. Table S6**). Because SelHap operates on haplotypes defined in genomic windows, we evaluated both large-scale structural variation and sequence-level novelty. Specifically, we compared 15 samples selected with SelHap to 15 breeder-selected cultivars using three complementary analyses: visual inspection of synteny, structural variant (SV) counts derived from pangenome graph analysis, and assessment of the increase in single-copy pangenome size.

A first visual inspection of SVs larger than 1 Mb suggests that the SelHap dataset contains more inversions and translocations than the breeders’ dataset (**Fig. 3A, B**). To quantify this trend across all SVs, we compared stepwise pangenome graphs constructed with Minigraph (8), and assessed robustness with 50 genome-order permutations. The SelHap dataset consistently showed a higher SV count, with a mean of 302,002 SVs, including 132 inversions, compared to 201,211 SVs and 60 inversions for the breeders’ dataset (**Fig. 3C**). Lastly, we analyzed the contribution of each dataset to the single-copy pangenome, which excludes repetitive regions using a k-mer-based approach, across the 53 domesticated samples of the BPGv2. The SelHap dataset presented a higher increase in single-copy sequences (**Fig. 3D**), indicating that it adds more novel sequence content to the barley pangenome. This result is consistent with the intended design of SelHap, which explicitly prioritizes conditional haplotype novelty relative to the existing pangenome background. Together, these analyses demonstrate that SelHap-based conditional novelty maximizes the addition of previously unrepresented sequence.

**Figure 3:**
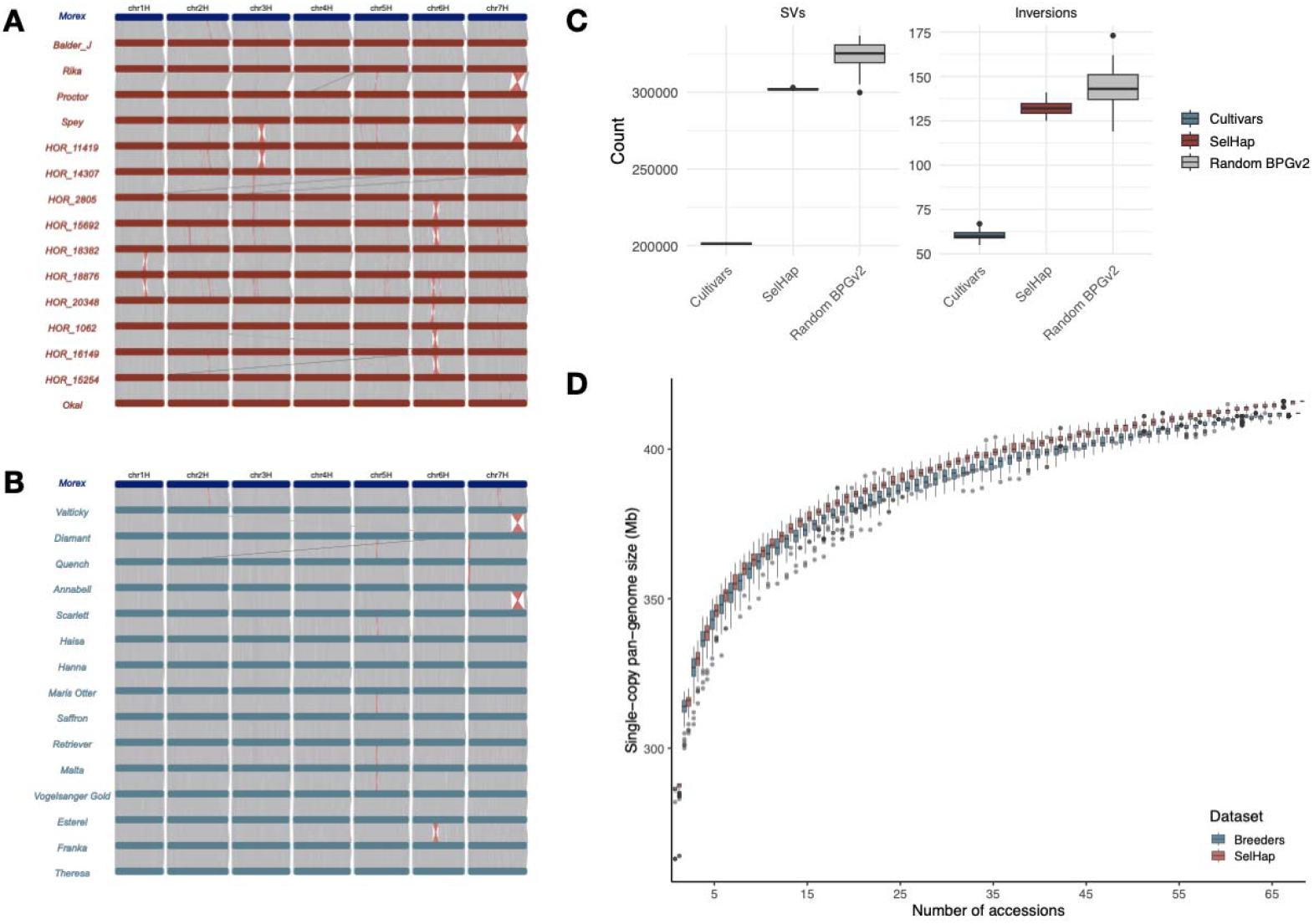
Comparison of SelHap dataset to the breeders’ dataset. **A, B:** NGenomeSyn (9) synteny plots of the SelHap dataset (A) and cultivar dataset (B). Gray bars represent syntenic regions between assemblies, red shows inversions and green translocations. Only alignments > 1000 kb are shown. **C:** Count of structural variants (SVs) and inversions from Minigraph analysis. Cultivars and SelHap datasets have 50 ordered permutations, whereas the BPGv2 (barley pangenome V2) dataset has 50 random subsamples of the total of cultivars and landraces from the BPGv2. **D:** Single-copy pangenome sizes of the two datasets and 53 domesticate samples from the BPGv2 (7), showing complexity estimated by single-copy k-mers. The curves trace the growth of non-redundant single-copy sequences as sample size increases. Error bars are derived from 100 ordered permutations each. The central line represents the median; the box bounds are the interquartile range (IQR), from the first quartile to the third quartile; and the whiskers extend to the most extreme values within 1.5⍰×⍰IQR from the quartiles

To control for potential confounding effects arising from differences in germplasm composition and sample selection strategy, we performed two additional analyses. Since sample order had a minor effect on Minigraph results (SD ± 338 and ± 466 for SelHap and breeders’ datasets, respectively), we generated an additional comparison dataset by performing 50 random samplings of 6 cultivars and 9 landraces from the barley pangenome (total of 17 cultivars and 46 landraces). We computed pangenome graphs for each subset and found a mean of 325,000 SVs, slightly higher than in the SelHap dataset (**Fig. 3C**). It is important to note that the BPGv2 dataset was used as background for the SelHap run, which may introduce some degree of bias in this comparison.

Because breeding gene pools differ in demographic history and genetic composition, comparisons across heterogeneous germplasm groups may be confounded. To minimize this effect, we performed an additional comparison restricted to a single agronomic gene pool: spring two-rowed cultivars. We compared two datasets, both with eight spring 2-rowed barley cultivars from the BPGv2 plus eight spring 2-rowed selected by the breeders or by SelHap (**Sup. Table S6**). Consistent with previous results, the SelHap dataset contributed to a greater increase in the size of the single-copy pangenome (**Fig. 4A**), despite similar numbers of SVs and inversions across datasets (**Fig. 4B**). We note that the use of the BPGv2 dataset as background for SelHap may favor the identification of haplotypes complementary to this specific reference, an effect that should be considered when interpreting absolute differences between datasets.

**Figure 4:**
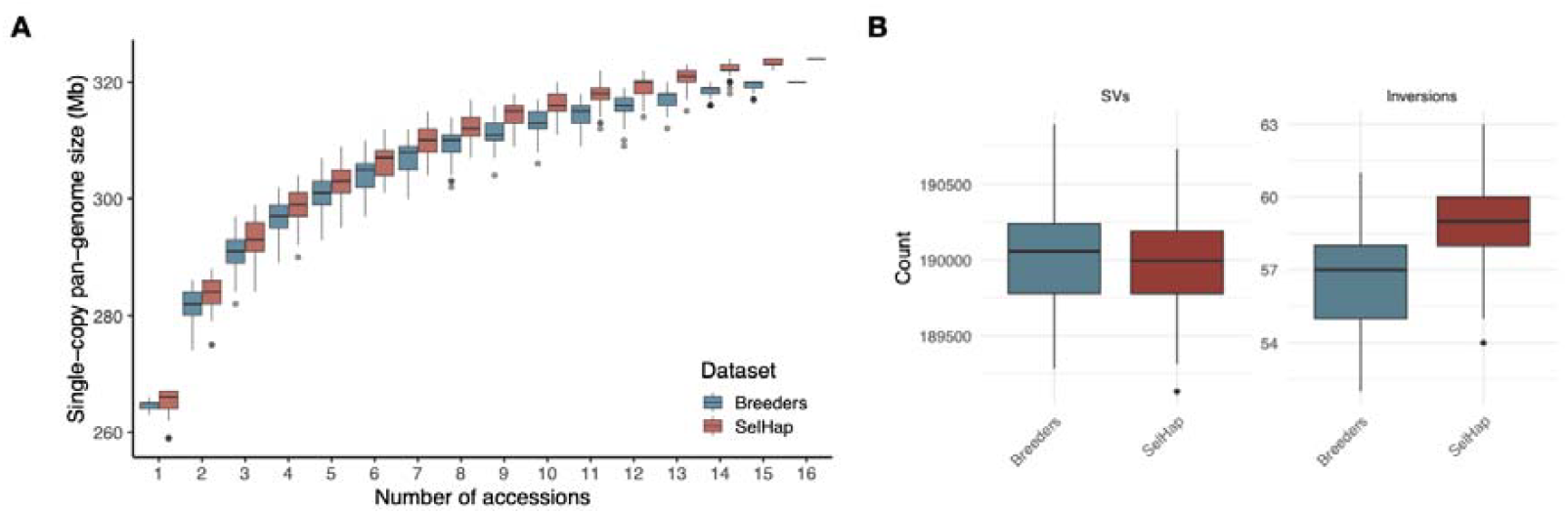
Comparison of two spring 2-rowed cultivars datasets. **A:** Single-copy pangenome sizes of the two spring 2-rowed cultivar datasets and 8 spring 2-rowed cultivars from the barley pangenome V2 (7) showing complexity estimated by single-copy k-mers. The curves trace the growth of non-redundant single-copy sequences as sample size increases. Error bars are derived from 100 ordered permutations each. The central line represents the median; the box bounds are the interquartile range (IQR), from the first quartile to the third quartile; and the whiskers extend to the most extreme values within 1.5⍰×⍰IQR from the quartiles. **B:** Count of structural variants (SVs) and inversions from Minigraph analysis with 50 ordered permutations each.

## Discussion

Using haplotype novelty relative to an existing pangenome background, SelHap consistently increased the addition of previously unrepresented sequence space. This effect was robust across diverse benchmarking scenarios, including comparisons to breeder-curated cultivar sets, random pangenome subsamples, and agronomically matched datasets. These results highlight a key shift in mature pangenome projects: once broad diversity is already represented, selection strategies must be conditioned on the existing pangenome composition to avoid redundancy and maximize information gain.

At earlier stages of pangenome construction, global diversity–based strategies (e.g., PCA on marker data) are appropriate and effective, and have successfully delivered a well-distributed representation of barley diversity in previous pangenome work (7, 10). As pangenomes approach saturation, the marginal gain of additional genomes decreases, and sample selection increasingly becomes a problem of avoiding redundancy relative to the existing pangenome composition. In such settings, SelHap provides a practical way to prioritize accessions that are most likely to contribute novel haplotypes relative to the existing background.

Rather than relying exclusively on global diversity summaries such as PCA or pairwise genetic distances, SelHap leverages haplotype information to identify genomic regions that are absent from an existing pangenome background. This focus on conditional novelty distinguishes SelHap from approaches that maximize overall diversity or feature counts without accounting for previously represented sequence. By summarizing haplotype-clustering output into interpretable scores and visualizations, SelHap enables targeted selection decisions that are directly aligned with the goal of extending mature pangenomes with genuinely novel sequence content.

Several practical considerations should be taken into account when applying SelHap. Sample rankings can vary with IntroBlocker parameter choices, including window size and SNP thresholds, which influence haplotype resolution and stringency. In heterogeneous panels comprising cultivars, landraces and wild accessions, multiple samples may exhibit comparable novelty scores, requiring additional judgment when prioritizing sequencing. Finally, SelHap relies on dense WGS data and is therefore less applicable to sparse marker datasets, although declining sequencing costs increasingly mitigate this limitation.

Together, our results demonstrate that conditioning sample selection on haplotype novelty relative to an existing pangenome provides a robust strategy for extending mature pangenomes. As pangenome resources continue to grow across plant species, approaches that explicitly target previously unrepresented sequence space will become increasingly important to maximize the return on sequencing and assembly efforts. SelHap offers a scalable and interpretable framework to support such targeted pangenome expansion in ongoing and future projects.

## Conclusions

As pangenomics advance, methods to complement already existing pangenomes without including redundant information become increasingly important. Here, we present SelHap and demonstrate its utility for identifying accessions carrying novel haplotypes for targeted pangenome expansion. Beyond sample selection, SelHap can support ancestry analyses, including comparisons between wild and domesticated germplasm or among different agronomic types. Together, our results demonstrate that conditioning sample selection on haplotype novelty relative to an existing pangenome provides an effective strategy for extending mature pangenomes while minimizing redundancy.

## Methods

### Data, short-read simulation, read mapping and variant calling

All data used is publicly available (**Sup. Table 1**). We used whole-genome shotgun (WGS) 150 bp paired-end short-read data of previously published panels: 210 2-rowed spring barley with around 4X coverage (11), and the barley 1000 core collection with coverages ranging from 3X to 10X (7), 315 modern cultivars with an average of 3X coverage (7), and 244 wild barleys with more than 10X coverage (6). We also used the 76 chromosome-scale barley genome sequences from the pangenome (7), comprising 17 elite cultivars, 36 landraces and 23 wild accessions. For these, we generated simulated paired-end reads with 10X coverage with fastq-generator v.1 (https://github.com/johanzi/fastq_generator) with parameters “generate_mapped_fastq_PE 150 0 10”.

Reads from all 1,845 samples were trimmed with Cutadapt v.1.15 (12) to remove adapters and subsequently mapped to the MorexV3 reference genome assembly using Minimap2 v. 2.24-r1122 (13). The resulting alignment files were sorted and duplicate-marked using Novosort v.3.0 and converted to CRAM files using SAMTools v.1.8 (14). SNPs were called using the ‘call’ and ‘mpileup’ functions of BCFTools v.1.15.1 (14) with minimum quality of 20. Variants were filtered based on read depth ratios using a custom awk script similar to the one at https://bitbucket.org/ipk_dg_public/vcf_filtering/src/master/ with the following parameters “--minmaf 0.05 --dphom 2 --dphet 1 --mindp 2000 --minhomn 1 -- minhomp 0.7 --tol 0.2 --minpresent 0.7 --minqual 40”.

### Obtaining clusters of haplotypes with IntroBlocker

We used IntroBlocker (3) to classify ancestral haplotype groups (AHGs) per 2Mb genomic window with a threshold of 500 SNPs per window and 400 SNPs for the Bayesian smoothing step. We defined the priority order based on pangenome, pangenome-wild, wild and domesticate accessions (**Sup. Table S7**). We modified the scripts to run each chromosome in parallel.

To assess sensitivity of SelHap to parameter choice, we performed additional analyses using different window size (5 Mb) and SNP thresholds (700 SNPs per window and 600 SNPs for Bayesian smoothing step). We ran SelHap on these datasets and compared the resulting scores and sample rankings across runs.

### Selection of samples with SelHap

We used the output tables after Bayesian smoothing as input for SelHap. These tables were loaded in R (function *read_tables()*) and formatted (*process_tables()*). We first ran the *compute_bin_haps()* function with the 76 pangenome samples as the background. This function assigns 0, 1 or 3 according to the absence, presence or missingness of a haplotype in a specific genomic bin. We ran it for 20 samples. To generate plots, we used the function *plot_results()*. The same process was repeated removing the wild samples from the panel (foreground), and from these results we selected 15 accessions for genome sequencing. We ran also *select_by_regions()* on the complete dataset to check for novel pericentromeric haplotypes. We used a range of +−80 Mb from the reported centromere regions in Morex (15) and selected 4 accessions from this list for genome sequencing.

### Selection of barley elite lines

Based on discussions with barley breeders, 13 elite cultivars were selected to be incorporated into the barley pangenome. We also searched through old breeding records (16), prioritizing accessions from before 1950 and directly derived from landraces across Europe. From those, we selected 6 that had seeds readily available.

### Genome sequencing and assembly

We obtained HiFi data (∼20-fold coverage) and Hi-C sequencing data (∼200M read pairs) for each genotype. High molecular weight (HMW) DNA isolation, library preparation and HiFi sequencing using the PacBio Revio device (Pacific Biosciences) were as described previously (17). Hi-C libraries were generated using the *DpnII* restriction enzyme as described previously (18). The libraries were sequenced (paired-end: 2 × 111 cycles) using a NovaSeq6000 device (Illumina) according to standard manufacturer’s protocols. Library construction and DNA sequencing were performed at IPK Gatersleben.

After obtaining the HiFi and Hi-C data, read quality control was performed using FastQC (19). PacBio HiFi reads were assembled using hifiasm v.0.24.0 (20). Pseudomolecules were constructed with the TRITEX pipeline (21) using Morex V3 (15) single-copy regions to guide scaffolding. Chimeric contigs and orientation errors were identified through manual inspection of Hi-C contact maps. Heterozygosity was assessed with Jellyfish v.2.2.6 (22) and GenomeScope v.2.0 (23). Genome completeness was evaluated using BUSCO (Benchmarking Universal Single-Copy Orthologs) v.3.0.2 (24) with embryophyta_odb10. In addition, assembly quality and completeness were further examined using Merqury (25), a k-mer-based approach that provides estimates of consensus quality (QV) and k-mer completeness independent of reference genomes.

### Gene annotation

Gene models were identified using a projection-based approach as previously described by Jayakodi et al. 2024. The underlying principles, detailed methodology, and source code of the projection pipeline are available at the GitHub repository (https://github.com/GeorgHaberer/gene_projection). Here, we describe only deviations in input data and parameter settings relative to previously published workflows.

High-confidence, evidence-based gene annotations of 20 *H. vulgare* and four diploid *H. bulbosum* accessions (FB19_011_3, FB20_005_1, FB20_029_7 and PI365428) were selected from previously published pan-genomic reports as starting source gene models (10, 26). Briefly, these 1,194,326 high-confidence input proteins were clustered at 100% sequence and size identity using CD-HIT (27), resulting in a non-redundant set of 559,842 barley transcripts and proteins. Functional classifications (protein-coding, plastid-related, and transposon-related models) were transferred from the corresponding source proteins. Orthologous groups were inferred using OrthoFinder v2.5.5 (28).

Protein and transcript sequences from the source models were aligned to the target genome using Miniprot (29) and Minimap2 (13), respectively. Only complete alignments featuring a contiguous open reading frame (ORF) and valid start and stop codons were retained for downstream processing.

The stepwise projection procedure comprised four sequential rounds: (i) selection of at most one protein-coding representative per orthogroup; (ii) inclusion of any remaining protein-coding models having a pfam hit with an e-value <10^−3^ ; (iii) projection of at most one plastid-related representative per orthogroup; and (iv) projection of at most one transposon-related representative per orthogroup. Completeness of the projected genes was assessed with BUSCO (24) employing the poales_odb10 reference set. Additional details on pipeline parameters and implementation are provided in the GitHub repository.

### Minigraph

Pangenome graphs were constructed with Minigraph. Because the graph is built stepwise, we did 50 permutations of order of genomes when constructing the graphs for each dataset. For the random BPGv2, we obtained 50 subsamples of 6 cultivars and 9 landraces to compute the graph. SVs were called for each repetition using gfatools bubble (v.0.5-r250-dirty, https://github.com/lh3/gfatools).

### Single-copy pangenome

Single-copy pangenomes were constructed as described previously (7) (https://bitbucket.org/ipk_dg_public/barley_pangenome/).

### Synteny plots

Whole-genome alignments of complete pseudomolecule assemblies were performed using Minimap2 v. 2.24-r1122 (13) with the −f 0.05 parameter to filter out repetitive minimizers. Visualization was done using NgenomeSyn (9).

## Availability of data and materials

The datasets generated during the current study are available in the ENA repository, under accession numbers PRJEB97413. The SelHap code is hosted in this repository: https://github.com/marimaro/SelHap with a step-by-step computational tutorial.

## Competing interests

The authors declare no competing interests.

## Funding

This research was supported by a grant from the German Ministry of Research, Technology and Space (BMFTR) to N.S., M.M. and M.S. (SHAPE-P3 [FKZ 031B1302A]).

## Authors’ contribution

M.M. and N.S. conceived and designed the study. M.P.M. and M.M. designed SelHap, and M.P.M. implemented the method. A.H. supervised genome sequencing activities. E.C. performed genome sequencing and contributed to genome assembly. E.C. and M.P.M. assembled the genomes. M.S. supervised and G.H. performed genome annotation. M.P.M. and M.M. wrote the manuscript with input from all coauthors.

## Acknowledgements

We acknowledge the technical assistance of Jacqueline Pohl, Sylvia Swetik, and Manuela Knauft during PacBio and Hi-C library preparation and DNA sequencing. We thank Anne Fiebig for sequence data submission. We thank Klaus Oldach for contribution with expert knowledge on barley elite cultivars.

## Notes

### Competing Interest Statement

The authors have declared no competing interest.

