## Supplementary_figures for "Selecting genomes that matter: haplotype-based prioritization for iterative pangenome expansion"

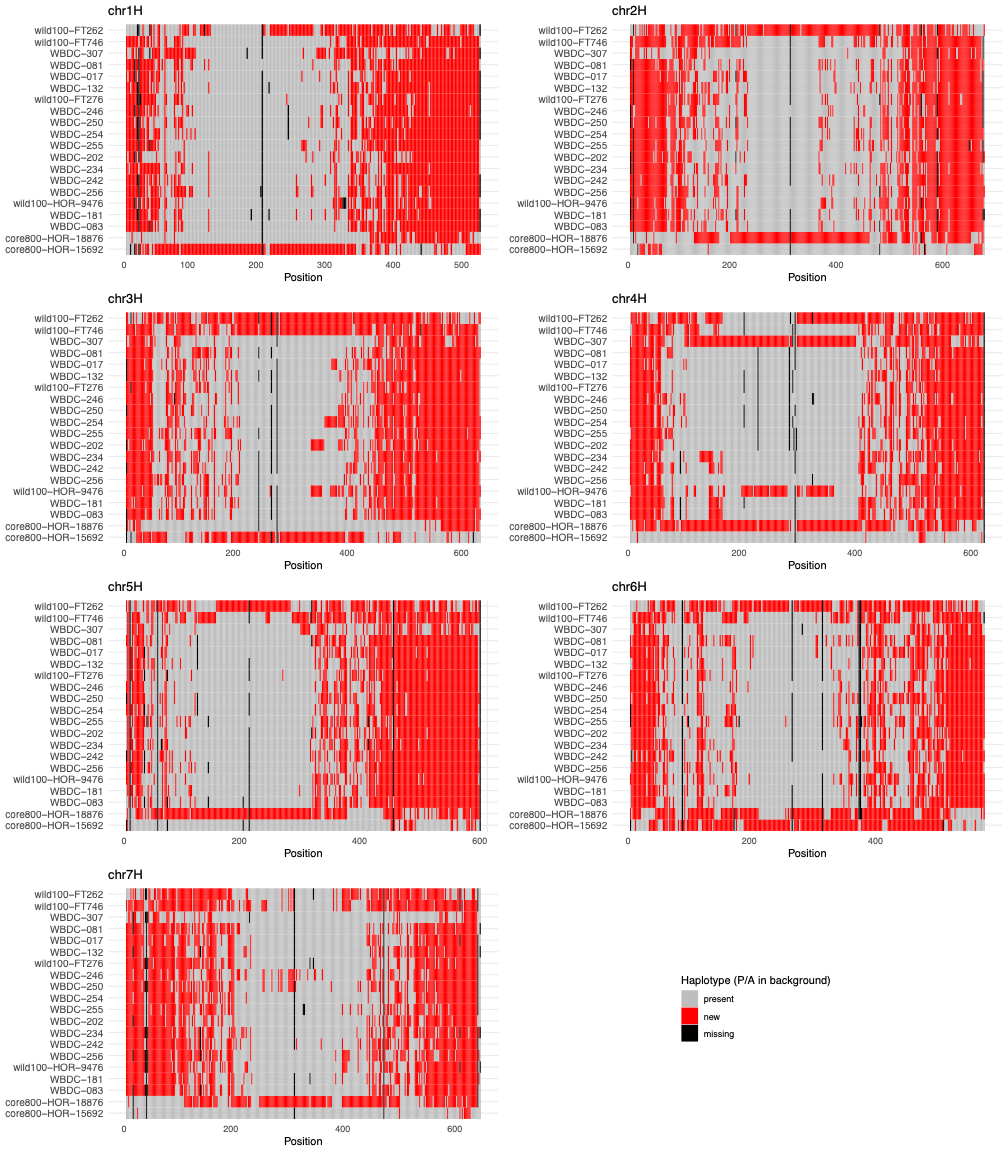

**Figure S1:** Binary haplotype plots showing novel (red) and background (grey) haplotype blocks across all chromosomes for the top 20 accessions selected by SelHap considering the whole dataset (1845 samples).

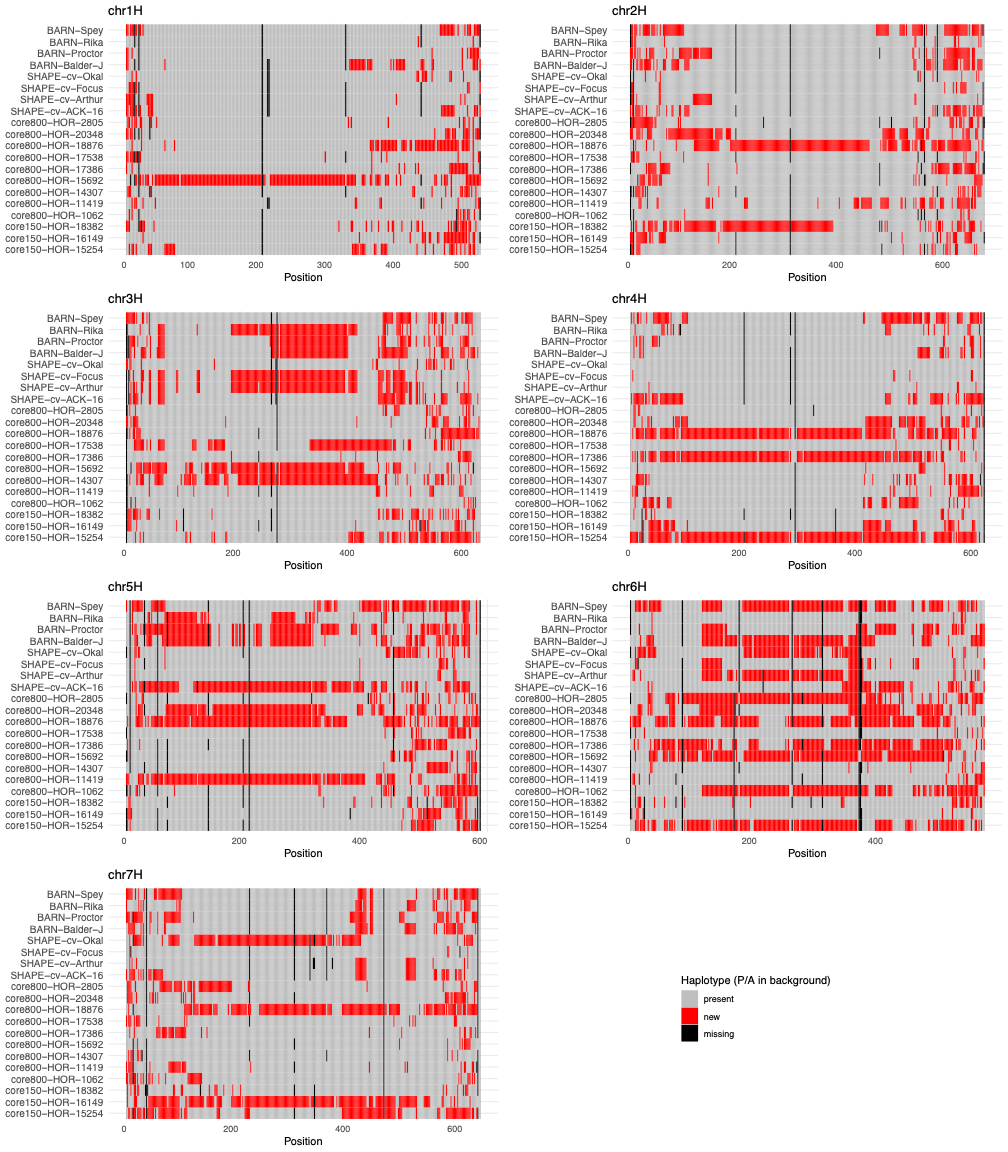

Figure S2: Binary haplotype plots showing novel (red) and background (grey) haplotype blocks across all chromosomes for the top 20 accessions selected by SelHap, excluding wild samples from the foreground (total of 1601 samples remaining). From these, 16 were selected to integrate the barley pangenome.

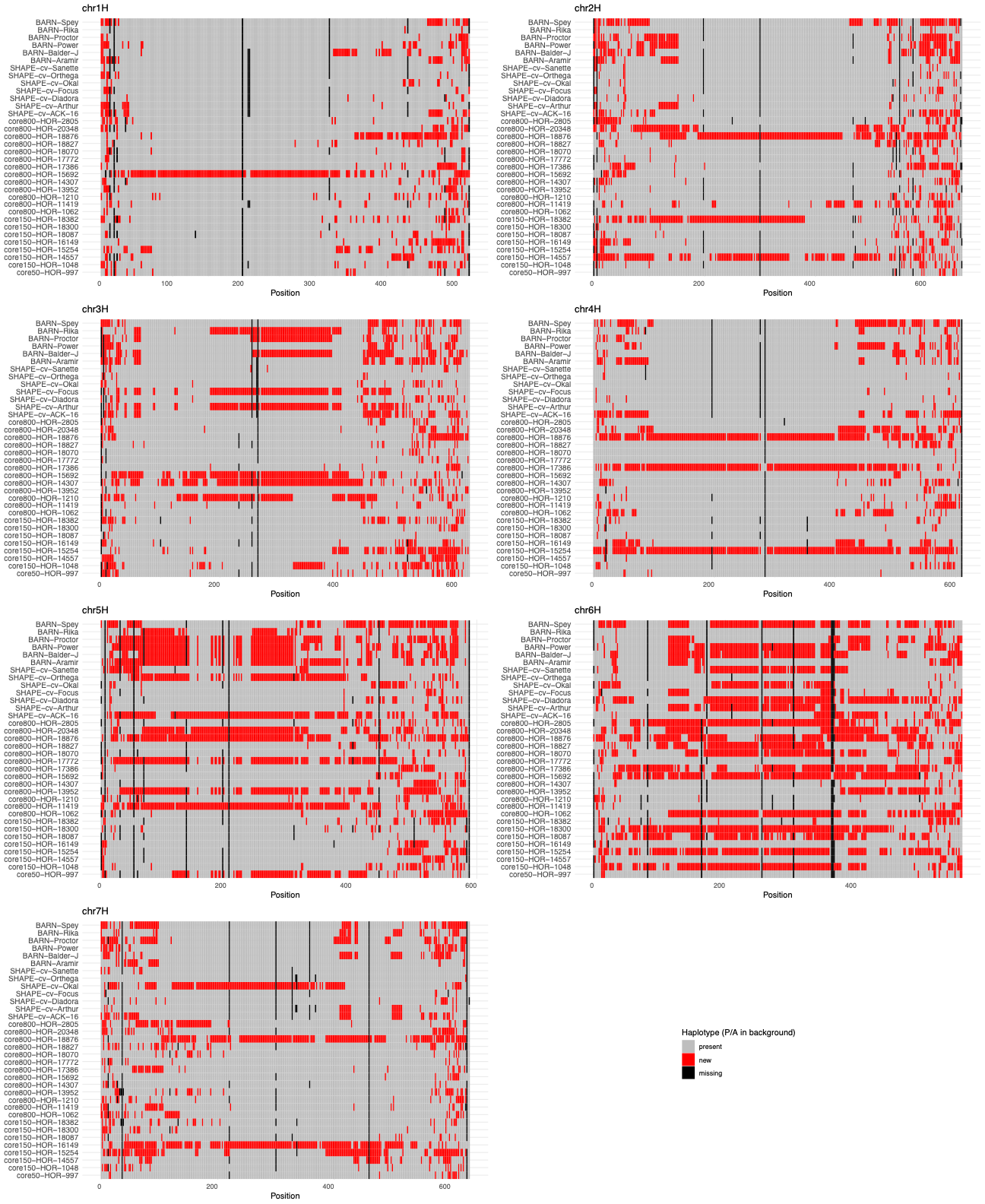

Figure S3: Binary haplotype plots showing novel (red) and background (grey) haplotype blocks across all chromosomes for accessions selected by SelHap with “select-by-region” mode for pericentromeric region (+- 80 Mb from MorexV3 centromere position). We selected 4 accessions from this list for genome sequencing.

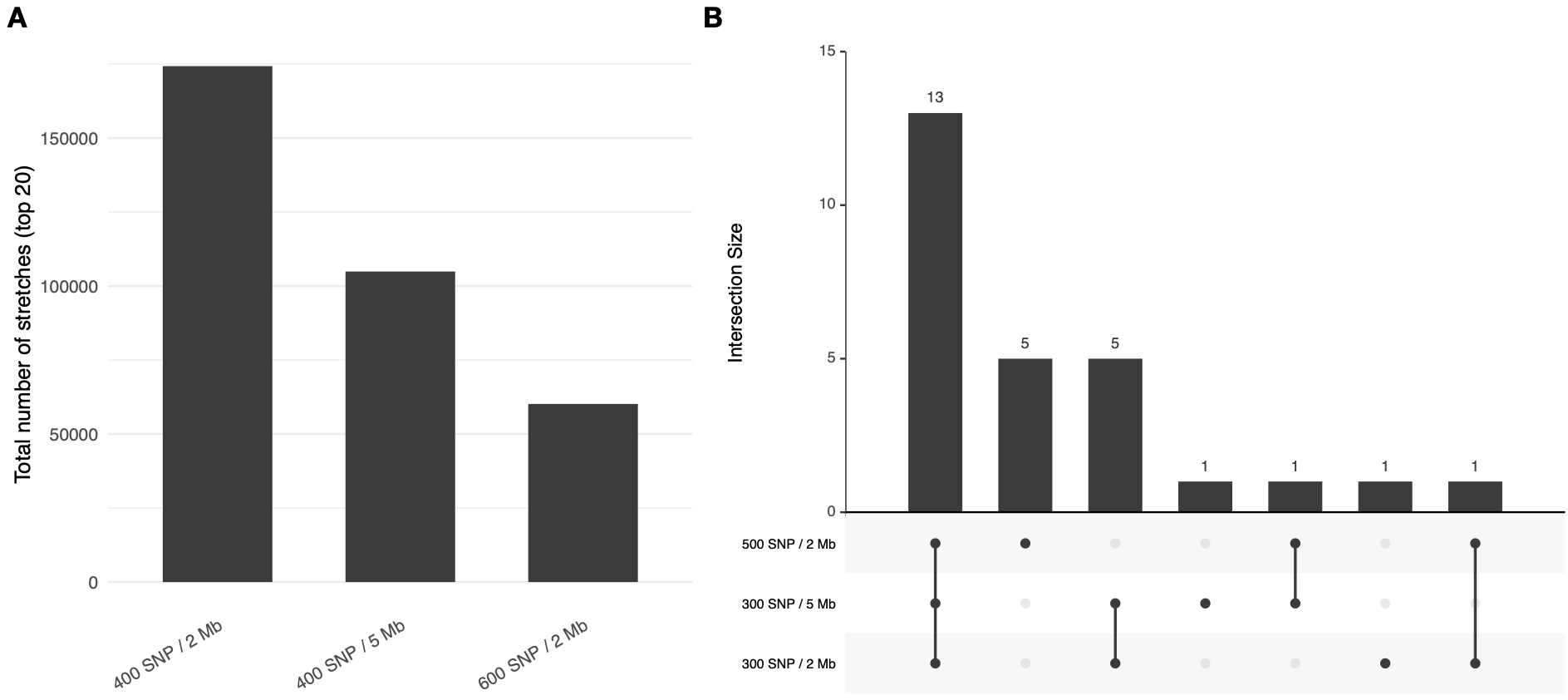

Figure S4: (A) Total number of novel haplotype regions contributed by the top-20 selected samples under the chosen parameters (300 SNPs threshold with 2 Mb window) and two alternative settings. Increasing window size or SNP threshold reduces detected novelty, consistent with increased stringency. (B) UpSet plot showing the overlap of top-20 selected samples across the 3 parameter settings.

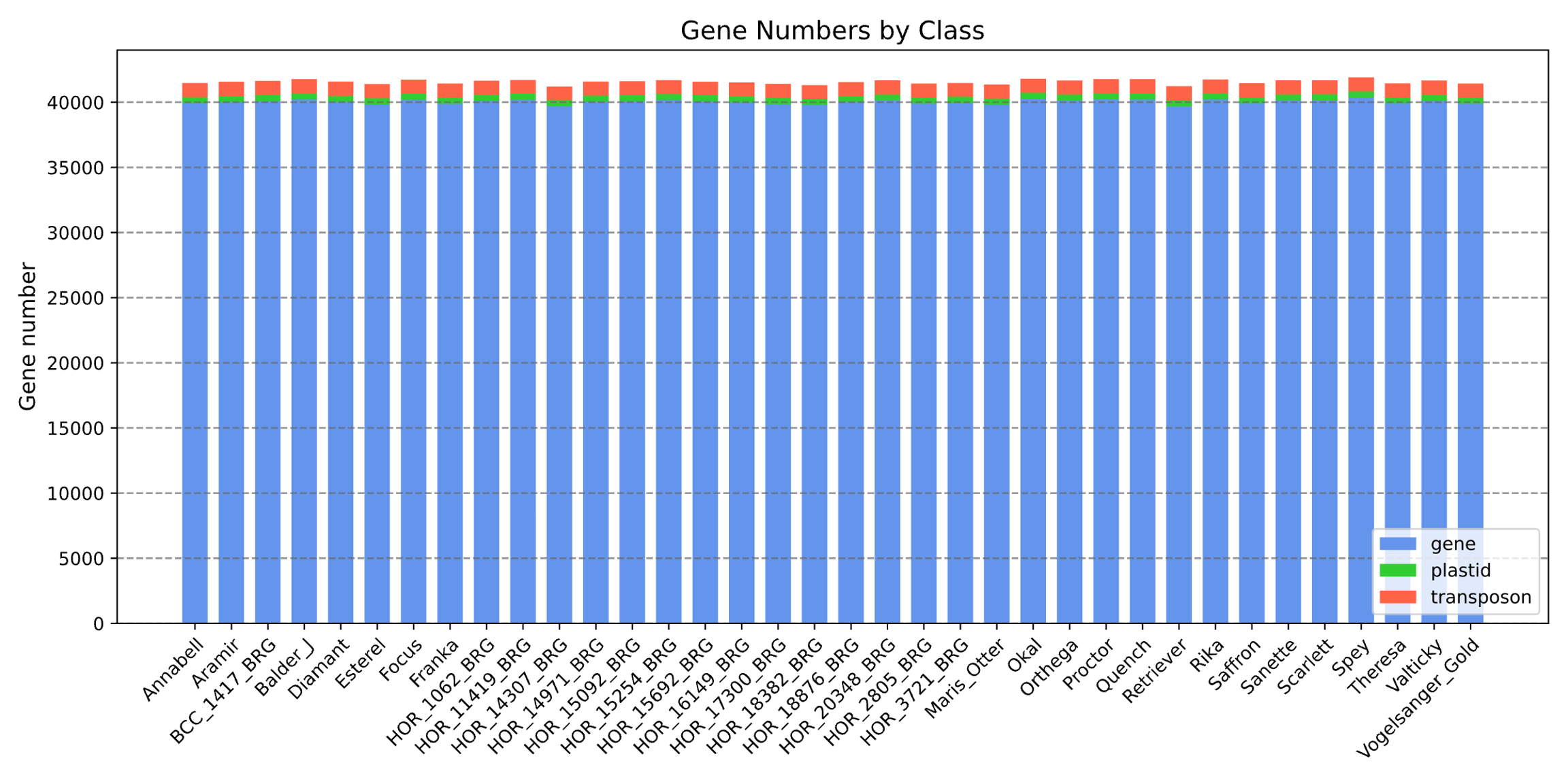

Fig. S5: Number of genes annotated per class in all sequenced accessions.

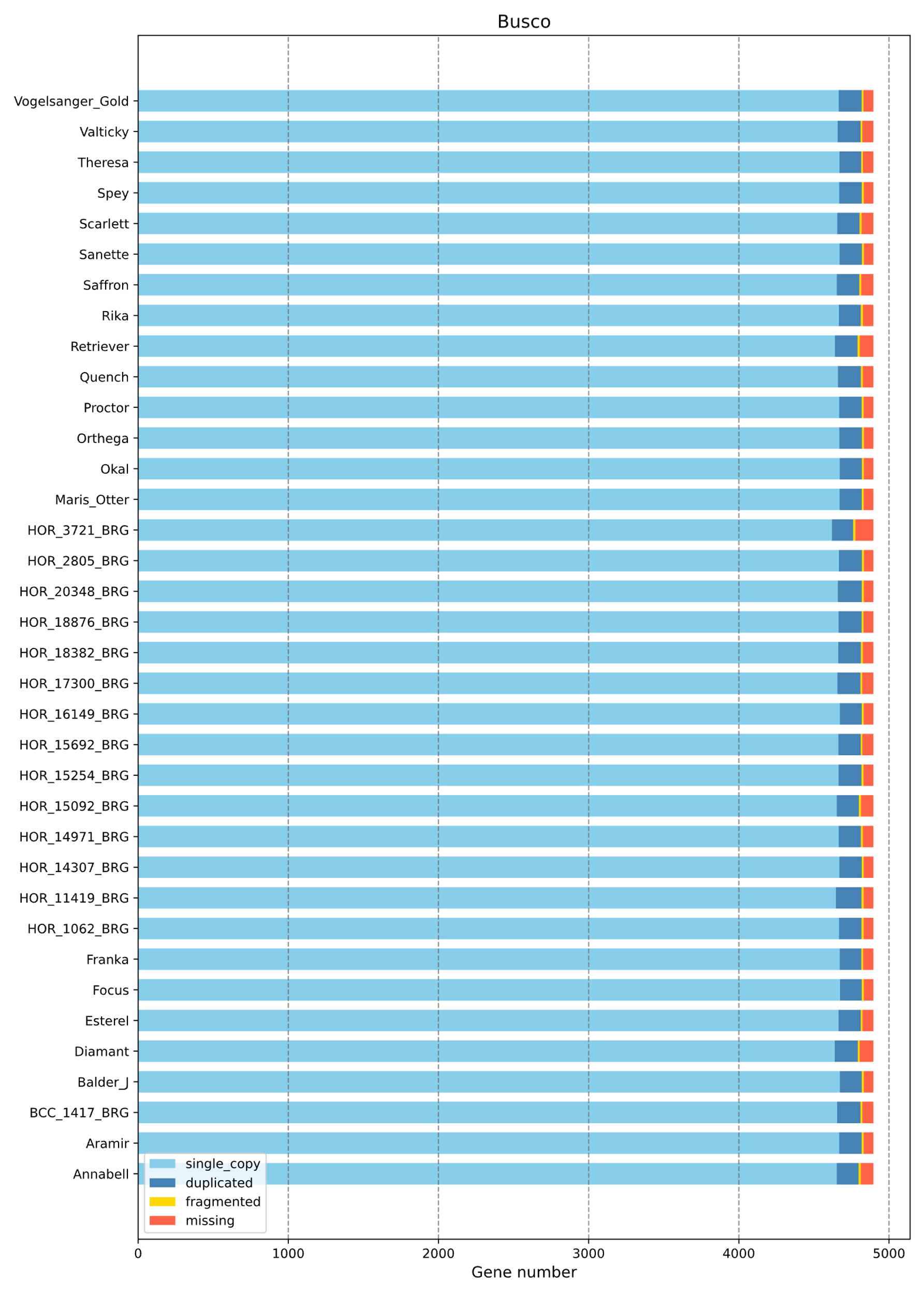

Fig. S6: BUSCO results after gene annotation for the 36 barley accessions using the poales-odb10 reference set.
